# Glioblastoma Drives Protease-Independent Extracellular Matrix Invasion of Microglia

**DOI:** 10.1101/2024.11.08.622715

**Authors:** Chia-Wen Chang, Ashwin Bale, Rohit Bhargava, Brendan A.C. Harley

## Abstract

Glioblastoma (GBM) is the most common and lethal form of primary brain cancer. Microglia infiltration into the tumor microenvironment is associated with immunosuppression and poor prognosis. Improved physicochemical understanding of microglia activation and invasion may provide novel GBM therapeutic strategies essential for improving long-term treatment efficacy. Here, we combine microfluidic systems with 3-D collagen hydrogels to systematically investigate microglia activation, invasion, contractility and cytokine secretion in response of GBM-microglia crosstalk. GBM inflammatory biomolecules significantly promote activation and 3D invasion of microglia. Interestingly, microglia invasion is not significantly affected by inhibitors of MMP activity or cellular glycolysis. In contrast, ROCK-pathway inhibition significantly impedes microglia invasion. Infrared microscopy analyses show that GBM co-culture does not significantly alter microglia lipid content. Further, GBM conditioned media resulted in significantly increased collagen hydrogel contraction, suggesting the importance of microglia contractility to physically remodel the local extracellular matrix (ECM). We also identify a panel of soluble proteins that may contribute to microglia chemotaxis, such as TIMP-1 and CXCL12. Taken together, this study suggests that the presence of GBM cells can enhance microglia invasion via increased cellular contractility, independent of MMP activity and cellular glycolysis.

## 1. Introduction

Glioblastoma (GBM), the most common and lethal form of primary brain cancer, has incredibly high mortality rates among all human cancers with a median survival of only 12 – 15 months.^[1]^ Despite an aggressive treatment strategy, including surgical resection followed by radiation and chemotherapy with Temozolomide (TMZ), long-term survival beyond 5 years is uncommon.^[2]^ Immunosuppression is one of the hallmarks of GBM that severely limits the efficacy of current therapeutic strategies, including immune-modulating treatments.^[3]^ The progression, therapeutic responses, and immunosuppression of GBM are also likely mediated by reciprocal interactions between GBM and its tumor-immune microenvironment (TIME). Multicellular tumor-immune interactions contribute to changes in the TIME landscape at the cellular, molecular and physical level which may influence GBM progression and therapeutic efficacy. Thus, there are urgent needs to improve our understanding of tumor-immune interactions.

The GBM TIME include multiple cell populations (e.g., GBM cancer cells, GBM stem cells, astrocytes, neurons, microglia and peripheral macrophages), vascular niches (e.g., pericytes and brain endothelial cells), and diverse extracellular matrix (ECM) components (e.g., collagen, hyaluronic acid).^[4]^ Microglia are the brain-resident immune cells that constantly survey their surroundings and maintain tissue homeostasis in the brain. Among all cellular components, microglia are the most abundant population of primary immune cells, accounting for up to 50% of GBM tumor mass.^[5]^ While many studies have extensively explored GBM cancer cell biology and underlying tumorigenesis mechanisms over last decades^[6]^, microglia have been largely underappreciated in GBM studies. However, emerging evidence suggests that GBM-microglia crosstalk may significantly modulate aspect of GBM progression: invasion, angiogenesis, therapeutic resistance, and a heightened immunosuppressive TIME.^[7]^ GBM cells may instruct microglia activation and infiltration, contributing to the abundant influx of microglia found within GBM tumors. Reciprocally, infiltrated microglia may also mediate GBM proliferation, migration, and therapeutic resistance via TIME remodeling or cytokine secretions. Targeting microglia via depletion, reprogramming, or altering recruitment kinetics, alone or combined with other therapeutic modalities, could reveal promising strategies to increase therapeutic efficacy. Recent progress in single-cell technology (e.g., single-cell RNA sequencing) has identified a variety of subtypes of tumor-associated microglia/macrophages in GBM.^[8]^ For instance, microglia are diffusely scattered throughout the GBM tumor but rarely present at the core of metastatic brain tumors, while peripheral macrophages often localize near CD31+ GBM vascular structures and throughout metastatic brain tumors.^[9]^ However, our understanding of dynamics of microglia invasion in GBM tumors is relatively scarce due in part to the intractable nature of deconstructing complex biochemical/cellular signaling and biophysical crosstalk *in vivo* as well as lack of effective models to recapitulate both cancer and immune components *in vitro*.

Tissue engineering approaches offer the opportunity to pattern multiple cellular constituents in tunable three-dimensional biomaterial models to reconstruct elements of the complex GBM TIME. We have previously developed gelatin and collagen hydrogels to explore the influence of aspects of the GBM TIME on GBM cell activity, including the role of matrix stiffness,^[10]^ hyaluronic acid content,^[11]^ and embedded microvascular networks^[12]^ on GBM cell proliferation, invasion, and drug response.^[13]^ Using these engineered models, we also showed GBM cell migration significantly decreased in response to co-culture with microglia.^[14]^ However, the reciprocal effect of GBM cells on the microglia behaviors, such as matrix invasion, has not been explored. Here, we used a 3-D microfluidic device to enable systematic evaluation of microglia matrix invasion, phenotype, and cytokine secretion profiles in responses to GBM cells (**Figure 1**). We report morphological changes of microglia in response to inflammatory biomolecules generated by GBM cells. We report the use of infrared spectroscopy to assess changes in microglia sub-cellular composition. We benchmark microglia matrix invasion in response to exogenous inflammatory biomolecules (e.g., LPS and IL-4), then define the influence of GBM conditioned medium as well as GBM co-culture on microglia invasion and microglia-mediated contractility. Furthermore, we characterize the cytokine profiles in the conditioned mediums collected from GBM, microglia and GBM-treated microglia culture. Collectively, our study provides novel insights of the biophysical dynamics of microglia matrix invasion in response to hydrogel models of the GBM TIME.

**Figure 1.**
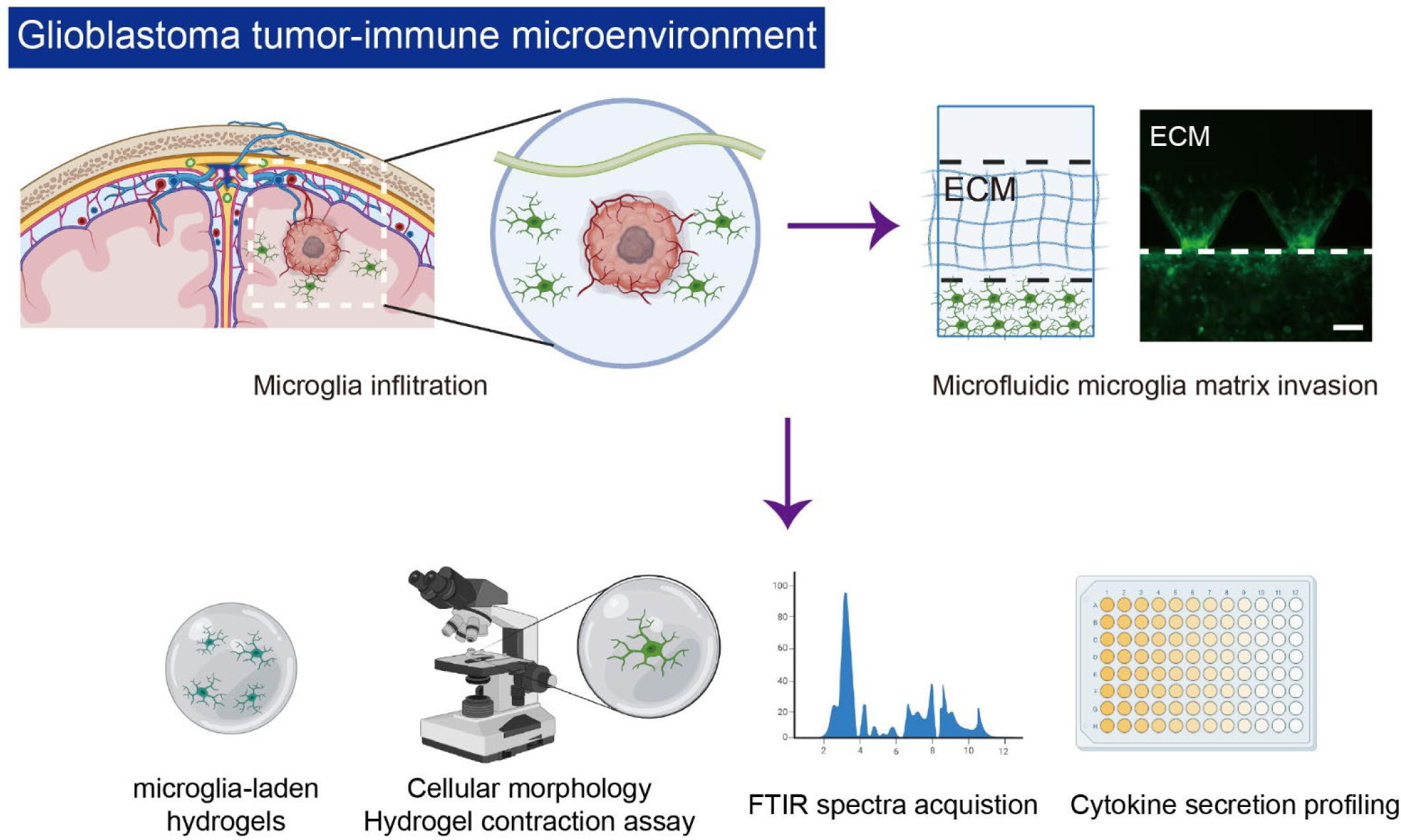
Schematic of tumor-immune microenvironment of glioblastoma (GBM). Highly infiltration of tumor-associated microglia is one of GBM hallmark. This study developed microfluidic models of GBM to study microglia matrix invasion. Cellular morphology of microglia cultured in 3D collagen hydrogels was observed to evaluate microglia activation in response to inflammatory biomolecules. Microglia-laden collagen hydrogels were used to evaluate the microglia cellular contractility by gel contraction assay. FT-IR spectroscopy was used to examine the biochemical fingerprints of microglia in various experimental conditions. The profiles of soluble cytokines and proteins were characterized using cytokine array approach. (This figure includes illustrations created by Biorender)

## 2. Results

### 2.1. Microglia exhibit altered morphology in response to GBM cells in collagen matrices

Microglial activation is associated with many central nervous system diseases, including brain cancers.^[15]^ Microglia undergo rapid morphological transitions, from ramified to hyper-ramified, during activation^[16]^ which involve elongation and increasing branching complexity.^[17]^ Shape index, a ratio of the perimeter to the area of an individual cell, can be used to quantify morphological changes of microglia (See “**Materials & Methods**”). Cell shape index of 1 represents a perfect round circle while the index of 0 refers to the indefinite elongated shape. To trace microglia activation in 3-D collagen hydrogels, we fabricated 3 mg/ml collagen hydrogels that have been previously used as a model the compliant brain matrix.^[18]^ Microglia cells were mixed with collagen hydrogel precursor solution, polymerized, then cultured (37°C) overnight prior to any biochemical treatments. Morphological changes to microglia were observed after 1 day culture via confocal microscopy (**Figure 1 and Figure 2**). While the average shape index of microglia is ∼ 0.57 for control cultures (**Figure 2A and 2D**), treatment with 50 ng/mL LPS as a positive control for inflammatory activation induced a significant decrease in shape index to ∼ 0.36 (**Figure 2B and 2D**). Exposure to GBM conditioned medium induced an even more significant decrease (mean shape index ∼ 0.25), suggesting GBM conditioned medium alone is potent enough to activate microglia (**Figure 2C and 2D**). These results demonstrate *in vitro* modulation of microglia activation is possible within a 3D collagen hydrogel via soluble inflammatory factors and that a range of responses to physiologic stimuli can be observed. These findings prompted us to subsequently evaluate microglia invasion potential in responses to inflammatory biomolecular signals.

**Figure 2.**
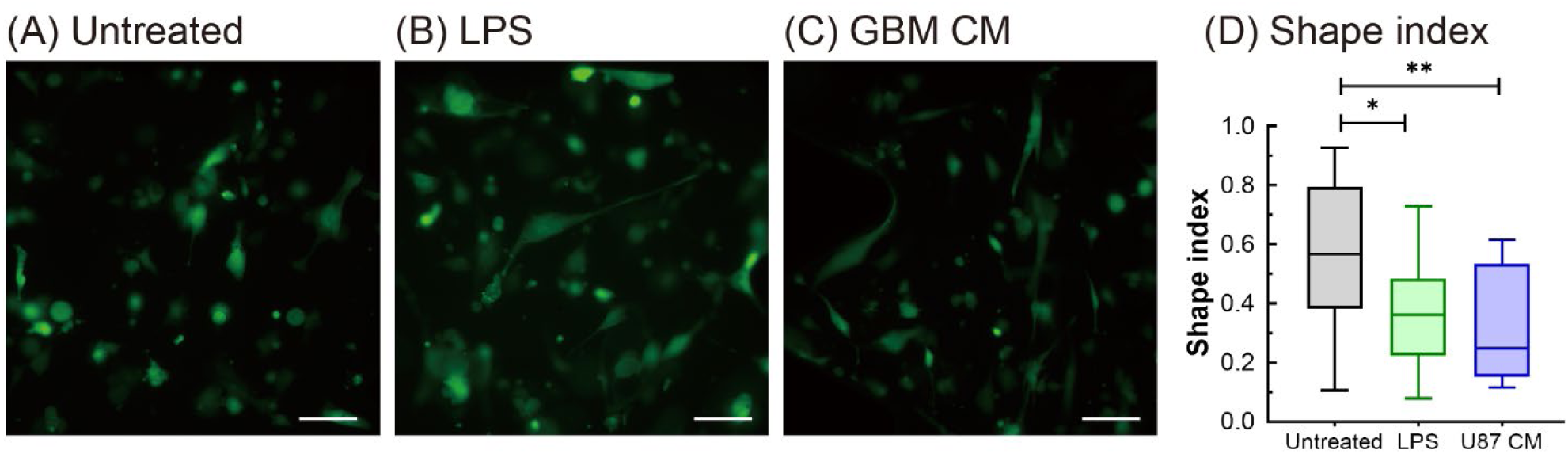
LPS and GBM conditioned medium treatments altered the microglia cellular morphology. Fluorescent images of microglia (green) cultured in 3-D collagen hydrogels in response to (A) untreated control, (B) LPS treatment and (C) GBM conditioned medium after 1 day culture. (D) Shape index analysis was applied to evaluate the morphological transition of microglia in various experimental conditions. The data were expressed as the box plot with quartiles (n ≥ 14). * and ** indicate a p value < 0.05 and < 0.01, respectively. Scale bars = 100 µm.

### 2.2. Inflammatory biomolecules and GBM crosstalk promote microglia activation and invasion

A 3-D biomimetic microfluidic system was used to study microglia invasion in response to biomolecular stimuli. The configuration of the microfluidic device was composed of a set of three parallel microchannels. The two outer channels were lined with human microglia cells (HMC3) while the central channel was filled with an acellular collagen hydrogel to form a stratified GBM TIME analogue (**Figure 1**). To assess the effects of microglia polarization on invasion, we compared microglia invasion in response to 50 ng/mL LPS and IL-4. LPS and IL-4 are known to activate microglia into pro-inflammation (or pro-M1) and anti-inflammation (pro-M2) phenotypes, respectively. Microglia showed minimal invasion into collagen hydrogels after 24 hours culture in the absence of LPS or IL-4 (**Figure 3A**). Interestingly, LPS-treated microglia show a 3.6-fold higher invaded area (increasing from 478 µm^2^ to 1735 μm^2^) compared to control (**Figure 3B and 3F**) cultures. Il-4 also increased the microglia invasion by 3.1-fold compared to control (from 478 µm^2^ to 1498 μm^2^) (**Figure 3C and 3F**). While both LPS and IL-4 treatments promoted significant microglia matrix invasion, there were no statistical differences between microglial invasion in responses to pro- vs. anti-inflammatory phenotypes.

**Figure 3.**
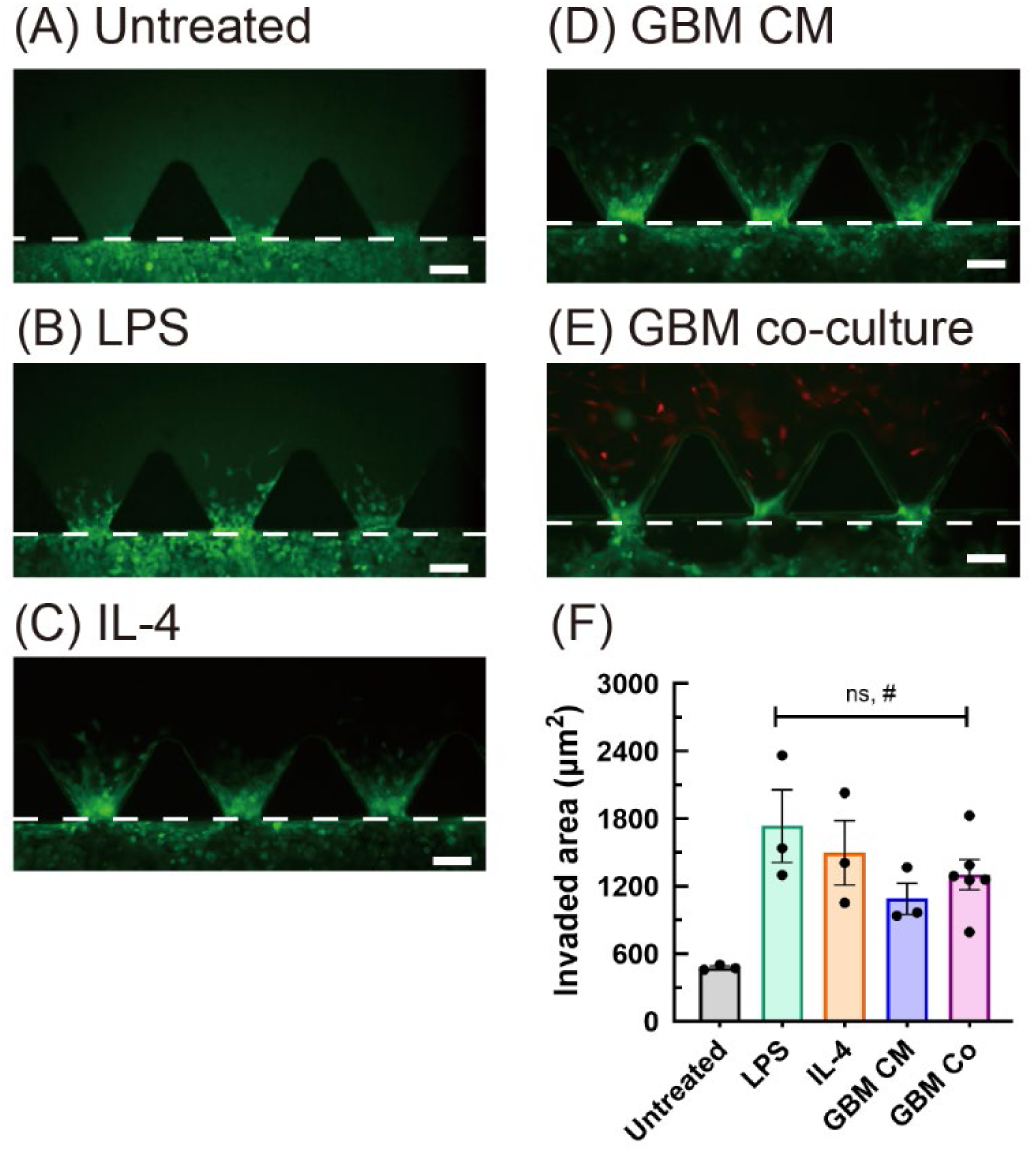
Microglia matrix invasion is promoted by various inflammatory biomolecules in GBM. Fluorescent images of microglia (HMC3, green) in the microfluidic device of (A) untreated control, (B) LPS, (C) IL-4, (D) GBM conditioned medium and (E) GBM co-culture conditions after 1 day culture. (F) averaged invaded area of microglia in the microfluidic devices. Microglia (HMC3, green) and GBM (U87, red) were used in GBM co-culture experiments. Dash white lines indicate the ECM/cell interfaces (Scale bars = 100 µm). The data were expressed as the mean ± standard error of the mean with individual data points (n ≥ 3 for all experimental conditions). ns indicates no significant difference and # indicates a p value < 0.05 compared to untreated control.

Next, we performed microglia invasion assays in a GBM-microglia coculture configuration where microglia were again seeded in the outer channels with the central channel now filled with U87-laden (1 x 10^6^ cells/mL) collagen hydrogels. Here, microglia invasion increased 2.7-fold compared to untreated control (478 µm^2^ to 1302 μm^2^) (**Figure 3E and 3F**). We hypothesized, given the short time of experiments, that microglia invasion was in response to paracrine signals from co-cultured GBM cells rather than local biophysical changes in U87-laden hydrogels. Thus, we measured microglia invasion in response to GBM conditioned medium. GBM conditioned medium also significantly promoted matrix invasion (1091 ± 139 μm^2^) with no statistical difference between microglia invasion in response to direct GBM co-culture (1302 ± 135 μm^2^) or GBM conditioned medium (**Figure 3D and 3F**). This suggests GBM cells could provide a chemotactic or chemokinetic effect to enhance microglia invasion. These results led us to choose GBM conditioned medium for further experiments to explore physicochemical mechanisms that mediate GBM-induced microglia invasion.

### 2.3. ROCK, but not MMP activity or cellular glycolysis, inhibit GBM-mediated microglia invasion

We subsequently assessed the role of matrix metalloproteinases (MMPs) on GBM-induced microglia invasion into collagen hydrogels. MMPs are known regulators of matrix degradation^[19]^ and are integral for angiogenic sprouting, vascularization, and cancer cell migration. ^[20]^ Nevertheless, our understanding of how MMP activity alters microglia movements are largely underdeveloped. We tested the effect of MMP inhibition using a full spectrum inhibitor of MMP activity (20 μM, GM6001, Millipore Sigma). Surprisingly, inhibition of MMP activity of microglia did not significantly influence GBM-induced microglia invasion into collagen matrices (881 ± 76 µm^2^) compared to untreated GBM conditioned media (1091 ± 139 µm^2^) (**Figure 4A**, **4B and 4E**). This interesting finding suggests that GBM-mediated microglia invasion may be largely independent of MMPs activity, prompting us to assess the roles of cell contractility (or cellular deformation) on GBM-induced microglia invasion. We hypothesized microglia may undergo rapid cellular deformation/contraction to navigate through the hydrogel matrix; we have previous reported the mean pore size within of these collagen matrices (∼ 1.7 μm), much smaller than the size of individual microglia.^[20a]^ Thus, we assessed microglia invasion in the presence of the Rho-associated, coiled-coil containing protein kinase (ROCK) inhibitor, Y27362 (20 μM, STEMCELL Technologies), a potent inhibitor of both ROCK1 and ROCK2 pathway. Notably, average GBM-induced microglia invasion (512 ± 68 µm^2^) decreased 53% and 42% in the presence of Y27362 compared to GBM conditioned medium (1091 ± 139 µm^2^) or GM6001-treated GBM CM conditions (881 ± 76 µm^2^), respectively (**Figure 4C and 4E**). GBM-treated microglia invasion in the presence of the ROCK inhibitor Y27362 (512 ± 68 µm^2^) showed statistically equivalent levels of invasion relative to untreated control (478 ± 13 µm^2^), suggesting GBM-promoted microglia invasion is primarily driven by cellular contraction.

**Figure 4.**
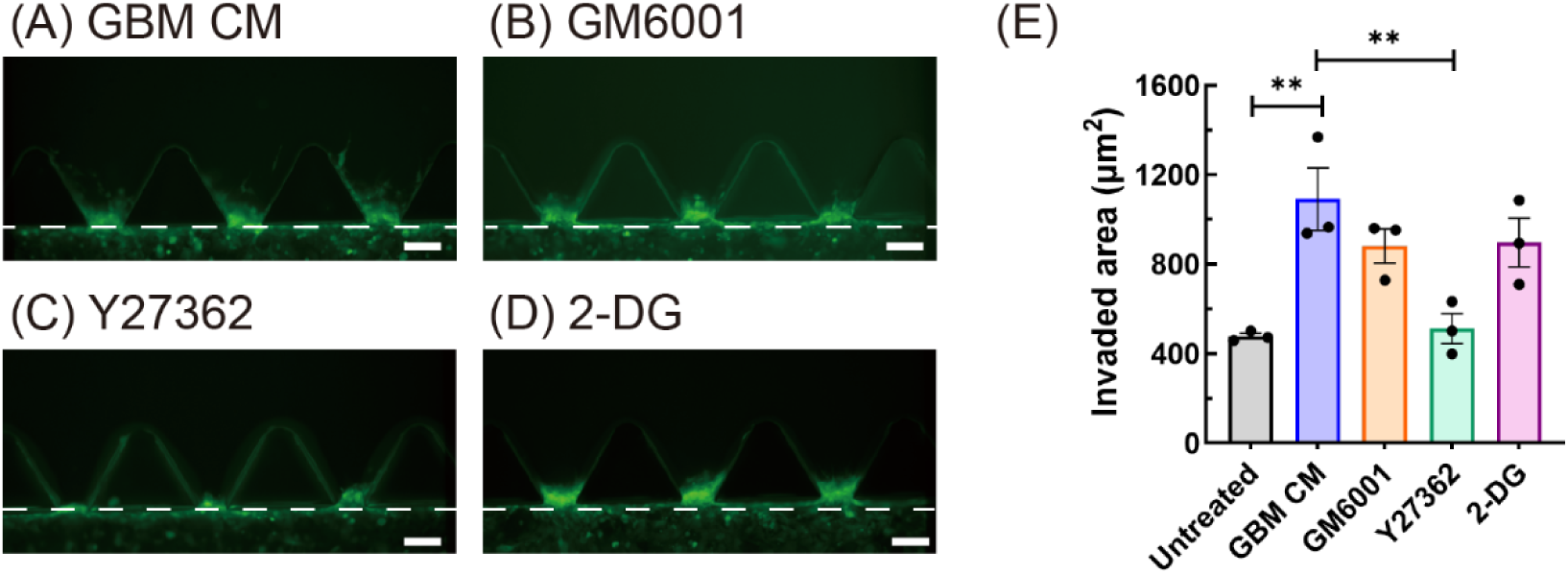
Microglia matrix invasion is mediated by various pharmacological inhibitors in the microfluidic devices. Fluorescent images of microglia invasion (HMC3, green) in responses of (A) untreated control, (B) GB6001 (C)Y27362 and (D) 2-DG treatments after 1 day culture in the devices. (E) average invaded area of microglia in the microfluidic devices. Dash white lines indicate the ECM/cell interfaces (Scale bars = 100 µm). The data were expressed as the mean ± standard error of the mean with individual data points (n = 3). ** indicates a p value < 0.01.

Metabolic reprogramming within tumor immune microenvironment, such as increasing cellular glycolysis, is known to influence cancer cell migration, metastasis, and therapeutic resistance.^[21]^ Further, activated microglia are known to switch their metabolic phenotype from oxidative phosphorylation (OXPHOS) to glycolysis for ATP production.^[22]^ Thus, it is possible that our observations of increased GBM-treated microglia invasion are also due to enhanced microglia cellular glycolysis. Using the same microfluidic device, we applied glycolysis inhibitor, 2-DG (1 mM), to test the roles of cellular glycolysis on microglia invasion in GBM. Here, average microglia invasion in the presence of 2-DG (897 ± 108 µm^2^) was not significant difference from GBM-treated control (1091 ± 139 µm^2^), suggesting shifts in cellular glycolysis was a not significant driver of microglia invasion in response to GBM (**Figure 4D**). Collectively, these results show that enhanced microglia invasion in response to GBM cells is driven primarily by the microglia contraction and that shifts in protease activity and cellular glycolysis are largely not involved in enhanced microglia invasion.

### 2.4. Microglia remodeled ECM by increasing gel contraction in response to GBM

Next, we measured the bulk contraction of microglia-laden collagen hydrogels as an indirect approach to quantify the cellular contractility and physical remodeling of microglia-laden collagen hydrogels in response to LPS, IL-4, or GBM conditioned medium. Here, a 1 mg/mL collagen precursor solution was mixing with microglia (1 x 10^6^ cells/mL) prior to any biochemical treatments. Microglia laden collagen hydrogels contracted 34 ± 3.9 % after 24 hours (untreated control), verifying their ability to contract ECM *in vitro* (**Figure 5A**). Microglia exhibited significantly increasd contractility in the presence of 50 ng/mL LPS (51 ± 4.8 % contraction) or 50 ng/mL IL-4 (49 ± 4.6 % contraction) compared to untreated control, respectively (**Figure 5**). Interestingly, GBM conditioned medium also significantly increased microglia-laden collagen gel contraction (46 ± 6.7 % contraction) compared to untreated control (**Figure 5D and 5E**). Hence, exposure to GBM conditioned medium increased bulk microglia contractility and microglia invasion driven by cellular contractility.

**Figure 5.**
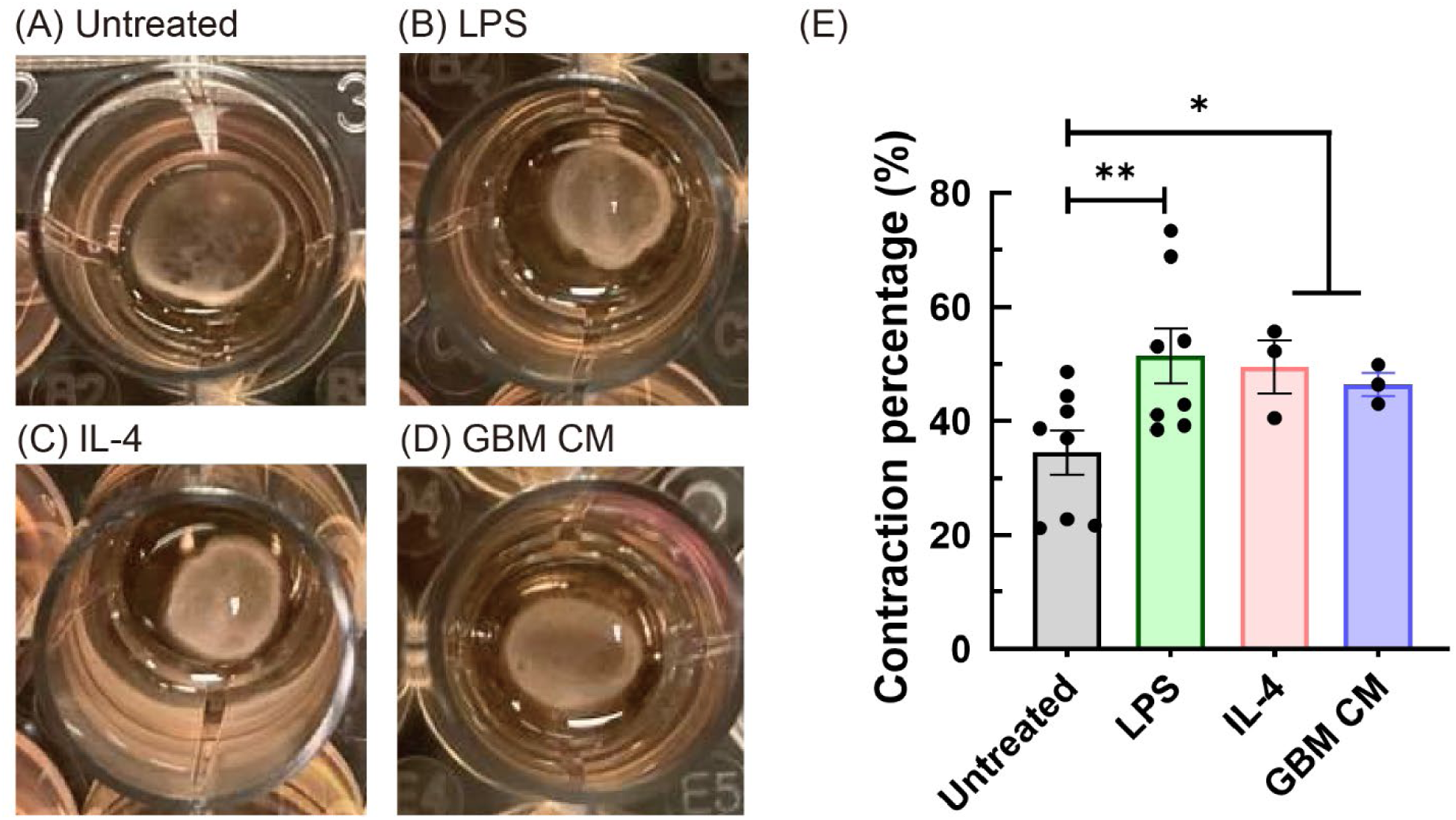
Gel contraction assay was used to evaluate microglia cellular contractility and its capabilities of physical ECM remodeling. Images of microglia-laden collagen hydrogels after 1 day culture in the conditions of (A) untreated, (B) LPS, (C) IL-4 and (D) GBM conditioned medium. (E) Contraction percentage of microglia-laden collagen hydrogels in various experimental conditions. The data were expressed as the mean ± standard error of the mean with individual data points (n ≥ 3). * and ** indicates a p value < 0.05 and < 0.01, respectively.

We subsequently used infrared (IR) spectral imaging^[23]^ to assess changes in the chemical status of microglia cultured in GBM CM or untreated control media, finding no significant difference across the spectral range (750-3800 cm^-1^) (**Figure 6A and 6C**). This data suggests microglia invasion is associated with microglia’s contractility rather than shifts in chemical processes, such as MMP proteolytic activities. Microglia treated with IL4 or LPS, however, greater cell-to-cell spectral variations (as indicated by the standard deviations in **Figure 6B and 6C**), indicating greater chemical heterogeneity within cells. These variations may be associated with different microglia polarization statuses (pro-M1 and pro-M2 phenotypes) in response to LPS and IL-4 treatments.

**Figure 6.**
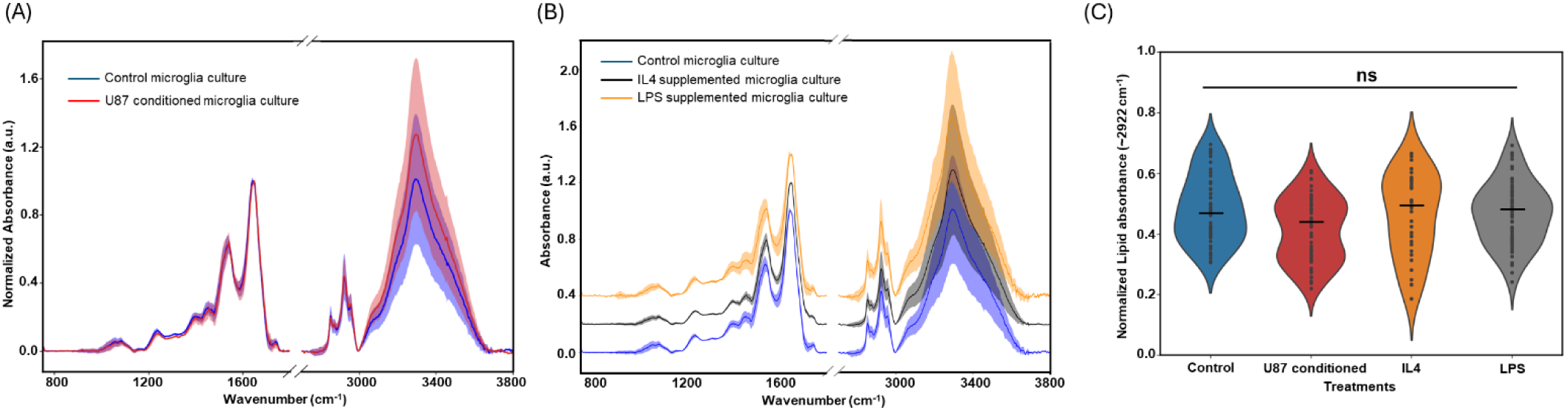
FT-IR spectroscopy was used to study the biochemical status of microglia in response to inflammatory biomolecules and GBM conditioned medium. (A) Normalized IR absorbance spectra from 750-3800 cm^-1^ wavelength with standard deviation marked as shaded area. (B) Normalized IR absorbance spectra from 750-3800 cm^-1^ wavelength averaged over 50-60 fixed cells, standard deviation marked as shaded area and spectra shifted vertically for visualization. (C) Lipid band (2922 cm^-1^) absorbance visualization for different treatment conditions. Each black dot represents 2922cm^-1^ band absorbance corresponding to individual cell. (n ≥ 50).

### 2.5. Cytokine profiling of GBM and microglia

We next sought to profile the composition of the conditioned mediums to identify potential proteins contributing to increased microglia invasion. We are particularly interested in distinguishing cytokine profiles of GBM and microglia cultures as well as the influence of GBM on subsequent microglia cytokine secretion. Thus, we examine cytokine expression in the following conditions using a cytokine array approach: 1) GBM conditioned media; 2) microglia conditioned media and 3) GBM-treated microglia conditioned media. We identified a set of proteins which were highly elevated in these conditioned media, mostly related to angiogenesis, ECM remodeling, inflammation, and chemokines (**Figure 7**, **8**). Considering first, angiogenesis and matrix remodeling-related cytokines, we observed increased expression of 13 proteins in U87 conditioned media, 18 in MG conditioned media, and 18 in GM-treated MG conditioned media out of 21 cytokines. Specifically, TIMP-1 was significantly elevated in all the CMs, suggesting its important role in regulating GBM TIME. Interestingly, cytokines highly expressed in GBM conditioned media are mostly related to angiogenesis (IGFBP-1, IGFBP-3, uPA, VEGF). Microglia secreted more cytokines related to inflammation (EGF, IL-8, Pentraxin 3, Thrombospondin-1). We also observed a cohort of cytokines expressed in GBM-treated microglia conditioned media consistent with the individual condition media (IL-8, uPa, VEGF) as well as some suggesting differential regulation in response to GBM-induced changes in MG activity (PIGF, EGF, SerpinEq, TIMP-1) (**Figure 7**).

**Figure 7.**
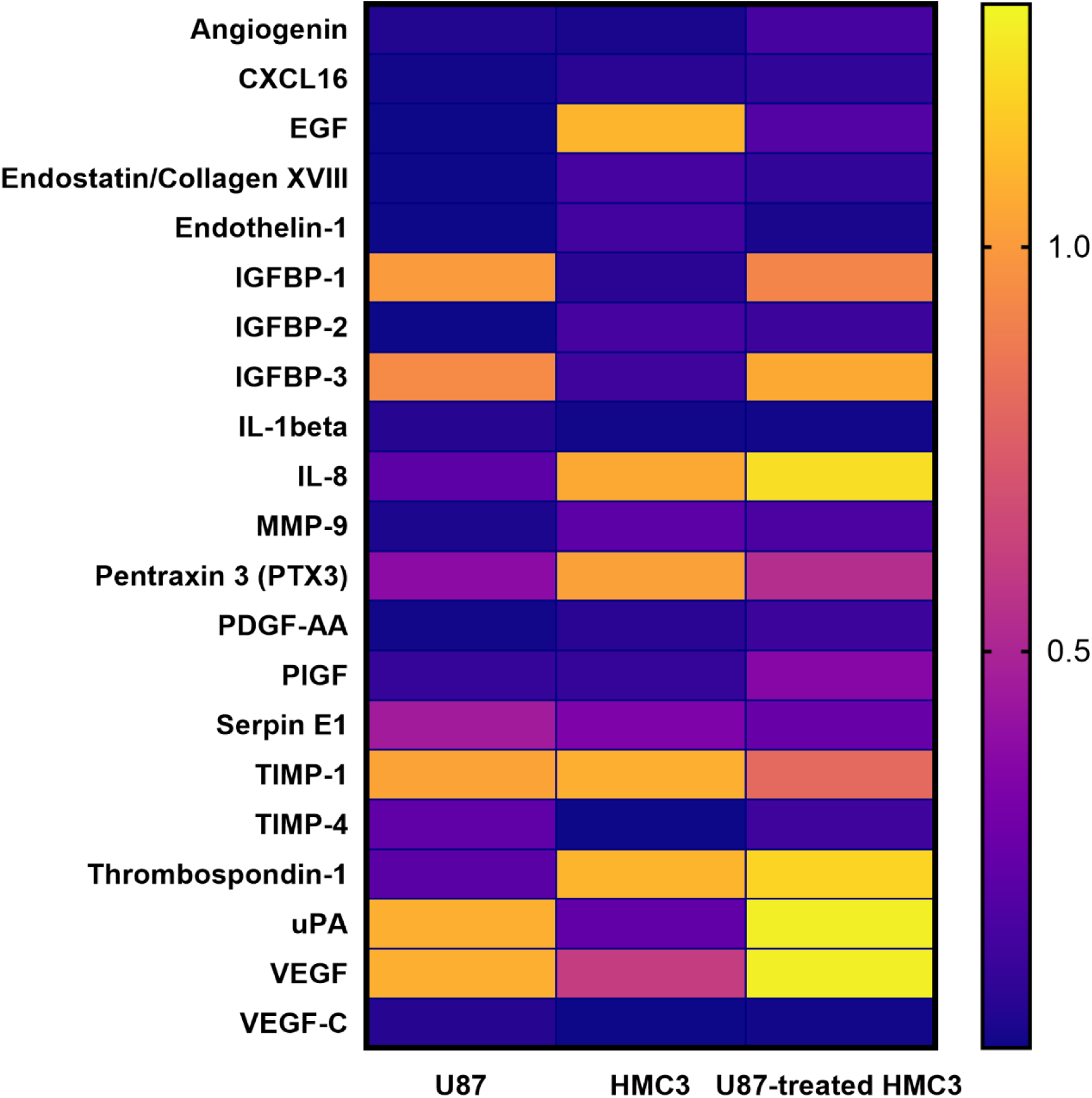
Expression of angiogenesis and matrix remodeling-related cytokines in the culture of GBM, microglia and GBM-treated microglia. The data was expressed as heat map which shows the normalized relative expression of 21 significantly detected matrix remodeling and angiogenesis-related cytokines in the secretome collected from U87, HMC3 and U87-treated HMC3 cells after 1 day of culture.

**Figure 8.**
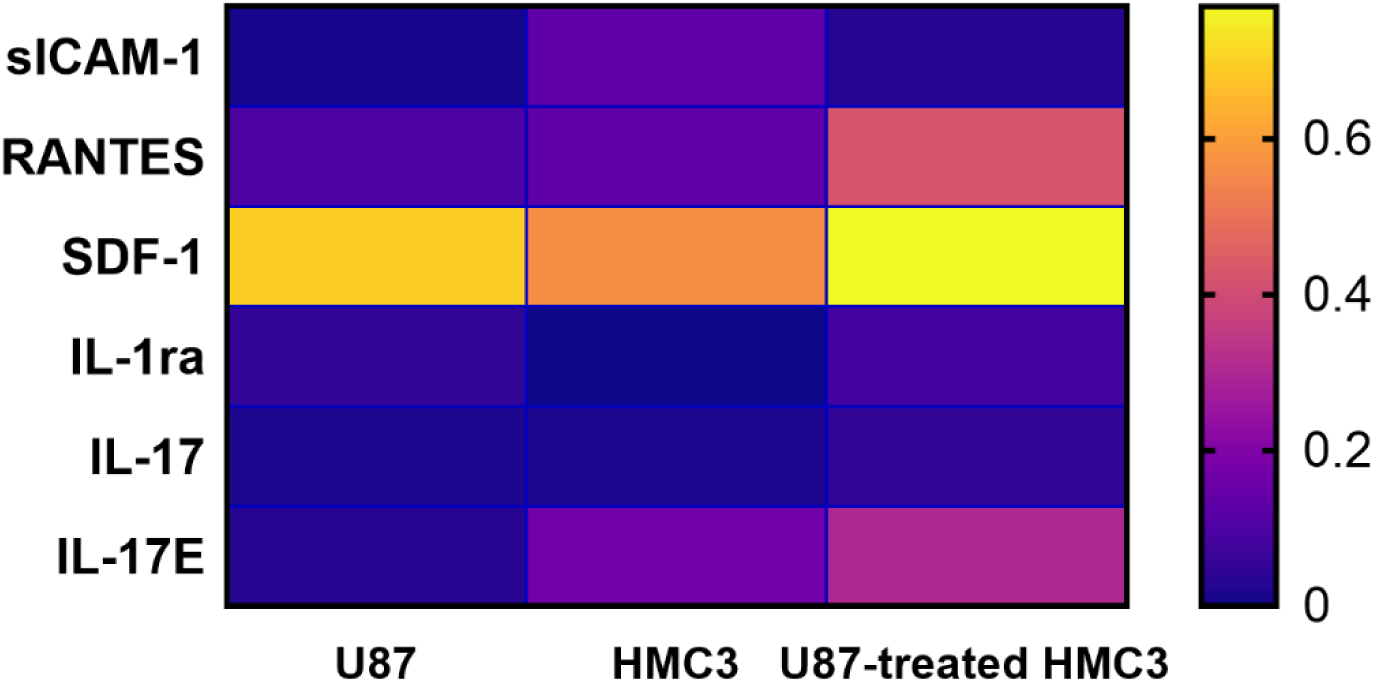
Expression of inflammatory-related cytokines and proteins in the culture of GBM, microglia and GBM-treated microglia. The data was expressed as heat map which shows the normalized relative expression of 6 significantly detected inflammatory-related cytokines in the secretome collected from U87, HMC3 and U87-treated HMC3 cells after 1 day of culture.

We then assessed expression level of inflammatory biomolecules in the same condition medias (**Figure 8**). We identified 6 inflammatory biomolecules (sICAM-1, RANTES, SDF-1, IL-1ra, IL-17 and IL-17E) which expressions were significantly increased (**Figure 8**). Importantly, we observed that the secretion levels of SDF-1 (a.k.a., CXCL12) were highly elevated across all condition medias. We also noticed that MG condition media shows significant higher secretion of cytokines related to pro-inflammation, such as sICAM-1, RANTES and IL-17E. Interestingly, the secretion levels of RANTES, SDF-1 and IL-17E appear to be increased in GBM-treated microglia condition media.

## 3. Discussion

Glioblastoma remains the most fatal and malignant form of primary brain cancer. Tumor-associated microglia and peripheral macrophages are the most abundant immune cells in the GBM TIME, accounting for up to 50% whole tumor mass. GBM cells are known to secrete an array of cytokines and proteins that may modulate microglia activation, recruitment, and invasion into tumor core. Microglia activation may also be influence by biophysical and biochemical properties of TIME (e.g. cellular components and ECM composition). Given the important roles of microglia in GBM progression, it is critical to identify processes that may regulate microglia invasion that may inform future therapeutic strategies.

Advancing our knowledge of microglia-GBM crosstalk requires a reproducible experimental system. While mice models, such as xenografts, are often used for drug studies, it is challenging to decouple physicochemical influences of GBM TIME on microglia activity *in vivo*. Currently, most *in vitro* microglia studies are performed either on 2D substrates or in Transwell inserts, oversimplified models of the brian ECM that do not provide a three-dimensional environment to assess invasion.^[24]^ Recently, 2D culture on hydrogel surfaces have been used to resolve the influence of T cells on microglia activation.^[25]^ Microglia have been shown to exhibit stiffness-dependent activation in response to inflammatory cues on 2D polyacrylamide gels.^[26]^ Two-dimensional uniaxial stretch platforms have also shown mechanical activation can promote microglia phagocytic and synaptic stripping activities.^[27]^ Nevertheless, these *in vitro* models are not readily amenable to conduct three-dimensional invasion assays. There is an opportunity to consider physicochemical features of native 3D ECM components in studies of microglia activation and invasion. In the present study, we fabricated tissue-engineered GBM models that incorporate 3-D ECM-derived hydrogels, strategies to spatially pattern microglia vs. GBM cells, and conditioned media approaches to study microglia invasion *in vitro*. This bioengineering approach allows us to consider multiple pathways that may regulate GBM-induced microglia invasion.

Here, we observed that microglia invasion significantly increased in response to both GBM cocultures or GBM conditioned medium. Interestingly, our finding shows that broad spectrum MMP inhibition (i.e., GM6001), nor inhibition of microglia cellular glycolysis (2-DG), did not reduce microglia invasion in response ot GBM conditioned media. In contrast, we found that microglia invasion significantly decreased when in response to ROCK-pathway inhibition (i.e., Y-27362) compared to GBM conditioned medium in the presence or absence of MMP and cellular glycolysis inhibition. Exposure to GBM conditioned medium also significantly increased microglia-mediated collagen hydrogel contraction verifying microglia contractility may play a significant role in physical remodeling of the GBM TIME. Collectively, our findings suggest that GBM enhance microglia invasion through increasing cellular contractility, independent of MMP activity and cellular glycolysis. While MT1-MMP expression by microglia has been recently associated with enhanced invasion of adjacent glioma cells,^[28]^ far less attention has been played on the processes that shape microglia motility. Improved understanding of spatial reorganization of microglia around the GBM microenvironment may offer new insight about shaping the neuroinflammatory environment of GBM.

Given that both MMP and glycolysis are chemically driven processes, the use of techniques like Fourier Transform Infrared (FTIR) spectroscopy can prove to be useful in providing unbiased full biochemical fingerprints of cells.^[[30]^ FTIR is a label-free non-destructive tool for studying molecular compositions and obtaining quantitative biomolecular measurements in terms of nucleic acids, carbohydrates, proteins and lipids in fixed cell specimens ^31]^. IR spectra measured for microglia cultured in both untreated control and GBM CM showed substantial overlap. This indicates similar biomolecular content in cells at the time of fixation for both culture conditions. MMP activity has previously been linked to membrane lipid raft formation to facilitate cell migration and invasion.^[32]^ Hence, our observation of minimal differences in the lipid band (∼2800-3000 cm^-1^) content of microglia in response to GBM conditioned media or control media supports the idea that changes in the microglia invasion is independent of MMP activity. Exogenous inflammatory LPS and IL-4 treatments, on the other hand, induced much greater spectral variations between cells. While these treatments have previously been known to induce pro-M1 (for LPS) and pro-M2 (for IL4)-like phenotypes in microglia ^[33]^, IR chemical imaging data suggests a wide spectrum of cell phonotypes may be present depending on their level of polarization.^[34]^ Indeed, it has been shown that microglia exhibit a multiple levels of polarization phenotypes.^[35]^ Notably, observed spectra variations for MG exposure to GBM condition media were much smaller than MG exposed to LPS or IL-4, suggesting the potent effects of GBM cells on microglia activation.

A major finding of this study is that MMP inhibition via GM6001 did not significantly alter microglia invasion, suggesting that microglia invasion is at least partially independent of MMP activity. This result is interesting because MMP is known to regulate several cellular activities in the brain tied to cancer cell migration and angiogenesis, notably that interstitial flow-mediated angiogenesis is partially dependent upon MMP activity.^[20a]^ MMP proteolytic activity has also shown to regulate the invasiveness of breast cancer cells.^[20c]^ This MMP independent invasion response seen in microglia is consistent with the trace secretion levels of MMP-9 in GBM and microglia culture, suggesting that in short term culture, GBM CM treatments did not significantly alter the microglia mediated matrix remodeling. Therefore, our findings suggest that microglia used cellular deformation and contractility driven by ROCK pathway to navigate through pores or spaces between collagen fibers. Indeed, increasing evidence also suggest that some immune cells, such as leukocytes and T cells, preferentially move through the submicron sizes holes or pores of tissues during the pathological progression.^[36]^ Since the progression of many inflammatory disease progression, including cancer, could potentially alter both ECM compositions and biophysical properties of tissue matrices, the findings from the present study will prompt future work to investigate the diseased ECM regulate immune cells dynamics. Separately, ROCK inhibition has been in involved in modulating microglia cytokine release and phagocytosis in neurodegenerative disorders and traumatic brain injury.^[37]^ Two ROCK inhibitors, Fasudil and Ripasudil, have already been licensed in Japan for clinical treatments of cerebral vasospasms and glaucoma.^[37b]^ Therefore, the findings of this study suggest a potential therapeutic target of microglia using ROCK inhibitors in GBM.

We observed distinct features of cytokine secretion when profiling the panels of 55 soluble factors secreted from GBM, microglia and GBM-treated microglia (See **Supplementary Information** for full list of soluble factors). The increased expressions of GBM-derived cytokines are mostly related angiogenesis while microglia promoted the secretion of cytokines related to inflammation. Interestingly, GBM CM significantly increased secretion of IL-8, PlGF, uPa and VEGF, demonstrating the essential roles of GBM-microglia crosstalk in shaping the biochemical landscape of GBM TIME. Notably, we noticed that secretion level of TIMP-1 is highly expressed both in microglia and GBM cultures, suggesting its important roles in mediating GBM progression. The overexpression of TIMP-1, a protein that involved in matrix degradation, is associated with tumor progression and poor prognosis in many cancers, such as pancreatic and breast cancer.^[38]^ Increasing evidence also suggests that its multifaced function in promoting immune cell infiltration in cancer and inflammatory diseases.^[39]^ In addition to TIMP-1, the secretion of SDF-1 (CXCL12) also showed significant increase in both GBM and microglia CM. CXCL12 is a key mediator of angiogenesis and metastasis in many tumors, such as breast cancer.^[40]^ In the context of GBM, CXCL12 promotes vasculogenesis, increases the resistance to radiotherapy and promotes tumor recurrence.^[41]^ Since the functions of TIMP-1 and CXCL12 are related to matrix properties and remodeling, it would be interesting for future work to define the interplay of ECM properties and these cytokines on orchestration of immune cell response (e.g., microglia) in brain cancer.

To our knowledge, this report is the first to describe biophysical processes used by microglia to invade into collagen matrices in responses to GBM cells. It is important to recognize that despite collagen not being the most abundant ECM in the normal brain, its content significantly increases in many brain pathologies, including GBM. Also, collagen I has been shown to promote tumorigenesis of glioblastoma stem cells through its binding with CD133.^[42]^ We note that GBMs secrete excessive hyaluronic acid (HA) during cancer progression and the accumulation of HA has been shown to promote GBM proliferation and migration. Therefore, it would be interesting in future study to investigate the role of HA on microglia activation and invasion in GBM. Collectively, we show that a combination of engineering approaches to study microglia invasion and GBM crosstalk can yield important insight into the role of the TIME in critical cancer processes.

## 4. Conclusions

Expanding the understanding of GBM-microglia interactions that contributes to microglia invasion into the GBM TIME provides new insight about future therapeutic strategies. Using 3-D microengineered collagen hydrogels embedded into a multi-channel microfluidic device, we investigated how GBM cells alter patterns of microglia invasion. GBM co-culture or conditioned media alters microglial activation but did not substantially alter their cellular composition. Presence of GBM cells significantly promoted microglia invasion into collagen hydrogels. Interestingly, increased microglia invasion is driven by cellular contractility and was independent of MMP activity or cellular glycolysis. We found TIMP-1 and CXCL12 were highly expressed in GBM, microglia, and GBM-treated microglia cultures. Take together, these findings refine our understanding of GBM-mediated microglia invasion. Furthermore, results from this study may inform future developments of therapeutics targeting microglia in the GBM TIME as well as introduce bioengineering tools to interrogate the complex dynamics of GBM-immune cell crosstalk in the tumor microenvironment.

## 5. Materials and Methods

### 5.1. Fabrication of microfluidic device

A microfluidic device with three sets of channels (AIM Bio) were employed to fabricate GBM TIME models for studying microglia invasion. The lateral channels were exploited for microglia cell culture and collagen-based hydrogel was introduced in the ECM compartment. Three channels were separated by series of trapezoid micropillars with a pitch of 200 µm. The width of openings (i.e., apertures) between the pillars were 100 µm to stably confine the hydrogel within the ECM compartment while allowing the microglia to contact with the ECM for matrix invasion. The widths of microglia channels are 500 μm. The ECM compartment was 1,300 µm wide and 250 μm deep. Type I collagen (Corning Inc.) isolated from rat tail was introduced into ECM compartment and polymerized at 37 °C in a humidified incubator overnight prior to microglia seeding into the adjacent media channels.

### 5.2. Preparation of GBM and microglia cultures

U87 (GBM) and HMC3 (microglia) cells were purchased from ATCC and are cultured in DMEM (Invitrogen) and EMEM (ATCC), respectively. U87 were cultured in T-75 flasks in a humidified incubator at 37 °C and 5% CO_2_, with the culture media changed twice a week. HMC3 were cultured in T-75 flasks in the same incubator with media changes every 3 days. Cell passages number of 2 – 10 were used in this study. Cells were harvested from a flask using 0.05% ethylenediaminetetraacetic acid (EDTA)–trypsin (Invitrogen) for 4 min, and the cell suspension was centrifuged at 950 rpm for 4 min. To acquire a monolayer of HMC3 lining the microglia channels, 10 µL of a 1 × 10^6^ cells/mL cell suspension was introduced to the channel via pipette. For all experiments, biomolecules were introduced after cells had been cultured in the microfluidic device for 1 day to ensure that treatment had no effect on initial cell attachment. To collect GBM conditioned medium, U87 cells (5 x 10^6^ cells/mL) were cultured in 6 wells plates with 2 mL DMEM for 1 day. The cell culture media was replaced with 2 mL EMEM and U87 cells were cultured for 1 day until media collection. GM6001 (CC10, Millipore Sigma) was diluted to 20 μM in culture medium and introduced to HMC3 cultured in the microfluidic device for the MMP inhibition experiments. For ROCK pathway inhibition, 20 μM Y-27362 was introduced with HMC3 cells in the microglia channels. 2-DG was diluted to 1 mM in culture media and added with microglia for cellular glycolysis inhibition.

### 5.3 Shape index analysis

Cell shape index was employed to measure the roundness of a particular cell. Bright field images were acquired using Leica DMI4000 microscope for analysis. Fiji ImageJ was used to quantify the area and perimeter of individual cell and shape index was defined by (4π × Area)/Perimeter^2^. For each experimental test condition, more than 14 cells were analyzed. Cell shape index of 1 represents a perfect round circle while the index of 0 refers to the indefinite elongated shape.^[43]^

### 5.4. Fourier transform infrared spectroscopy (FT-IR) spectroscopic characterization of cell cultures

The pipeline for cell culture and FT-IR image acquisition has been illustrated in Figure S1. Cells corresponding to various treatment conditions were cultured on low emissivity glass slides prior to IR imaging. Cells were fixed in 4% paraformaldehyde for 20 minutes at room temperature, followed by three 1 x phosphate-buffered saline (PBS) washes. Samples were stored in a dry box until imaging and FTIR imaging was performed based on previously reported protocol.^[44]^ A Perkin Elmer Spotlight 400 (Perkin Elmer) spectrometer equipped with a highly sensitive linear MCT detector array was used. Images were acquired in reflection mode in the midinfrared range 750-3800 cm^-1^ at a spectral resolution of 4cm^-1^ and with 120 co-additions for 6.25 μm pixel size at line-scan of 2.2 cm/s. CellProfiler was used to segment cells.^[45]^ The resulting individual cell spectra were baseline corrected and normalized with respect to Amide I peak (∼1650 cm^-1^) and average IR spectra across multiple cells per condition were used to compare the visible differences between the different experimental groups.

### 5.5. Microglia invasion analysis

To visualize microglia matrix invasion, HMC3 cells were stained with CellTracker Green (5 μM, Invitrogen) for 45 minutes before cells were seeding into the microfluidic device. Leica DMi8 confocal microscope (Leica, 612 Germany) was used to capture microglia invasion from ECM/cell interfaces before (day 0) and after 24 hours of treatments (day 1) in all conditions, and FIJI imageJ was used to quantity the invaded area of each aperture (i.e., cell/ECM interface). At least 18 apertures were quantified in each microfluidic device and at least 3 microfluidic devices were analyzed as experimental replicates to perform statistical analysis.

### 5.6. Hydrogel contraction assay

The contractility of microglia was measured by collagen hydrogel contraction assay using previous reported approach.^[46]^ Collagen gels were prepared by adding microglia suspension to a 1 mg/ml type I collagen solution to make a final cell density of 1 × 10^6^ cells/ml. 150 µl of collagen solution with cells was cast into each well of a 48-well plate with at least three replicates per condition. Subsequently, collagen gels polymerized for 45 min at 37°C and 5% CO_2_ in a humidified incubator. Following 45 minutes of polymerization, an additional 200 µl of fresh cell culture medium was added to each well. For LPS and IL-4 treatments, LPS and IL-4 were reconstituted to a concentration of 50 ng/mL in fresh EMEM to passively diffuse into collagen hydrogels. For GBM treatments, 200 µL of U87 conditioned medium was added into each hydrogel. The surface area of the gels was quantified by FIJI ImageJ software at each time point. The percent contraction was determined by equation 1:

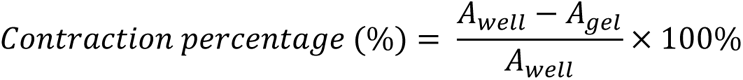

### 5.7 Cytokine profiling of GBM and microglia culture

A Proteome Profiler Human Angiogenesis Array (ARY007, R&D Systems) was used to determine relative levels of 55 soluble biomolecules related to angiogenesis and ECM-remodeling. Medium from the GBM, microglia and GBM-treated microglia cultures was collected at days 2 of culture in the tissue 6-well culture plate, stored at -80°C until use, and pooled for analysis. 500 μL of individual medium was used to cytokine profiling experiments. The array was run as per the manufacturer’s instructions and imaged using ImageQuant 800. Cytokine dot intensities were quantified using the MATLAB Protein Array Tool (https://www.mathworks.com/matlabcentral/fileexchange/35128-protein-array-tool). The negative control spot averages were subtracted from the pixel density of each sample, and then the pixel density of each sample was normalized to the pixel density of positive control spots. For inflammatory cytokines profiling, a Proteome Profiler Human Cytokine Array (ARY005B, R&D Systems) was used to determine relative levels of 36 proteins related to inflammatory signals. See **Table S1** and **Table S2** for full list of proteins assessed in the cytokine array experiments (**Supplementary Information**).

### 5.8. Statistical analysis

Numerical data reported in this manuscript were expressed as the mean ± standard error of the mean (SEM). Each experimental condition was performed at least in three replicates to conduct statistical analysis unless noted otherwise. Variations of all data were statistically analyzed by performing one-way analysis of Variance (ANOVA) followed by a post hoc unpaired, two-tailed Student’s t test, executed by OriginLab and GraphPad Prism software. To compare the statistical difference of each experimental condition, the asterisk mark (*) was applied as * for a p value of <0.05, ** for a p value of <0.01, and *** for a p value of <0.001.

## Conflicts of Interest

The authors declare that they have no competing interests.

## Acknowledgements

The authors would like to acknowledge funding from the National Cancer Institute under Award Numbers R01 CA256481 (BACH) and R01 CA260830 (RB). The authors also acknowledge additional funding provided by the Department of Chemical and Biomolecular Engineering, the Carl R. Woese Institute for Genomic Biology, and the Cancer Center at Illinois at the University of Illinois Urbana-Champaign. Chia-Wen Chang gratefully acknowledges the funding from IGB Fellowship Award at the Carl R. Woese Institute for Genomic Biology, University of Illinois Urbana-Champaign. The authors would like to acknowledge the following institutes for access to their facilities and services: the Carl R. Woese Institute for Genomic Biology and the Beckman Institute for Advanced Science and Technology, located at the University of Illinois. The authors thank Jung Bale for the assistance with preliminary data collection and analysis. The content herein is solely the responsibility of the authors and does not necessarily represent the official views of the National Institutes of Health.

## Contributions (CRediT: Contributor Roles Taxonomy ^[47]^)

**Chia-Wen Chang:** Conceptualization, Data curation, Formal Analysis, Visualization, Investigation, Methodology, Writing – original draft, and Writing – review & editing.

**Ashwin Bale**: Data curation, Investigation, Methodology, and Writing – review & editing.

**Rohit Bhargava**: Resources and Writing – review & editing.

**Brendan Harley:** Conceptualization, Resources, Project administration, Funding acquisition, Supervision, and Writing – review & editing.

## Notes

### Competing Interest Statement

The authors have declared no competing interest.

## References

[1] A. C. Tan, D. M. Ashley, G. Y. López, M. Malinzak, H. S. Friedman, M. Khasraw, CA: A Cancer Journal for Clinicians 2020, 70, 299.

[2] L. R. Schaff, I. K. Mellinghoff, JAMA 2023, 329, 574.

[3] a)B. T. Himes, P. A. Geiger, K. Ayasoufi, A. G. Bhargav, D. A. Brown, I. F. Parney, Front Oncol 2021, 11, 770561; b)M. S. Graham, I. K. Mellinghoff, Cancer Cell 2021, 39, 304.

[4] A. Bikfalvi, C. A. da Costa, T. Avril, J.-V. Barnier, L. Bauchet, L. Brisson, P. F. Cartron, H. Castel, E. Chevet, H. Chneiweiss, A. Clavreul, B. Constantin, V. Coronas, T. Daubon, M. Dontenwill, F. Ducray, N. Entz-Werlé, D. Figarella-Branger, I. Fournier, J.-S. Frenel, M. Gabut, T. Galli, J. Gavard, G. Huberfeld, J.-P. Hugnot, A. Idbaih, M.-P. Junier, T. Mathivet, P. Menei, D. Meyronet, C. Mirjolet, F. Morin, J. Mosser, E. C.-J. Moyal, V. Rousseau, M. Salzet, M. Sanson, G. Seano, E. Tabouret, A. Tchoghandjian, L. Turchi, F. M. Vallette, S. Vats, M. Verreault, T. Virolle, Trends in Cancer 2023, 9, 9.

[5] a)F. Khan, L. Pang, M. Dunterman, M. S. Lesniak, A. B. Heimberger, P. Chen, The Journal of Clinical Investigation 2023, 133; b)G. Wang, K. Zhong, Z. Wang, Z. Zhang, X. Tang, A. Tong, L. Zhou, Frontiers in Immunology 2022, 13.

[6] W. Wu, J. L. Klockow, M. Zhang, F. Lafortune, E. Chang, L. Jin, Y. Wu, H. E. Daldrup-Link, Pharmacological Research 2021, 171, 105780.

[7] H. Liu, Y. Sun, Q. Zhang, W. Jin, R. E. Gordon, Y. Zhang, J. Wang, C. Sun, Z. J. Wang, X. Qi, J. Zhang, B. Huang, Q. Gui, H. Yuan, L. Chen, X. Ma, C. Fang, Y.-q. Liu, X. Yu, S. Feng, Cell Reports 2021, 36, 109718.

[8] a)W. Wang, T. Li, Y. Cheng, F. Li, S. Qi, M. Mao, J. Wu, Q. Liu, X. Zhang, X. Li, L. Zhang, H. Qi, L. Yang, K. Yang, Z. He, S. Ding, Z. Qin, Y. Yang, X. Yang, C. Luo, Y. Guo, C. Wang, X. Liu, L. Zhou, Y. Liu, W. Kong, J. Miao, S. Ye, M. Luo, L. An, L. Wang, L. Che, Q. Niu, Q. Ma, X. Zhang, Z. Zhang, R. Hu, H. Feng, Y.-F. Ping, X.-W. Bian, Y. Shi, Cancer Cell 2024, 42, 815; b)A. T. Yeo, S. Rawal, B. Delcuze, A. Christofides, A. Atayde, L. Strauss, L. Balaj, V. A. Rogers, E. J. Uhlmann, H. Varma, B. S. Carter, V. A. Boussiotis, A. Charest, Nature Immunology 2022, 23, 971; c)P. Schmassmann, J. Roux, S. Dettling, S. Hogan, T. Shekarian, T. A. Martins, M.-F. Ritz, S. Herter, M. Bacac, G. Hutter, eLife 2023, 12, RP92678.

[9] a)W. Xuan, M. S. Lesniak, C. D. James, A. B. Heimberger, P. Chen, Trends in Immunology 2021, 42, 280; b)E. Friebel, K. Kapolou, S. Unger, N. G. Núñez, S. Utz, E. J. Rushing, L. Regli, M. Weller, M. Greter, S. Tugues, M. C. Neidert, B. Becher, Cell 2020, 181, 1626.

[10] a)J.-W. Chen, S. Pedron, B. A. C. Harley, Macromol Biosci 2017, 17, 1700018, 1700018; b)S. Pedron, E. Becka, B. A. C. Harley, Adv Mater 2015, 27, 1567; c)S. Pedron, B. A. C. Harley, J Biomed Mater Res Pt A 2013, 101, 3405.

[11] a)S. Pedron, G. L. Wolter, J.-W. E. Chen, S. Laken, J. N. Sarkaria, B. A. C. Harley, BioRxiv 2019; b)J.-W. E. Chen, S. Pedron, P. Shyu, Y. Hu, J. N. Sarkaria, B. A. C. Harley, Front Mater 2018, 5.

[12] a)M. T. Ngo, J. N. Sarkaria, B. A. C. Harley, Advanced Science 2022, 9, 2201888; b)M. T. Ngo, E. Karvelis, B. A. C. Harley, Integr Biol (Camb) 2020, 12, 139.

[13] V. A. Kriuchkovskaia, E. K. Eames, R. B. Riggins, B. A. C. Harley, Advanced healthcare materials 2024, e2400779.

[14] J.-W. E. Chen, J. Lumibao, S. Leary, J. N. Sarkaria, A. J. Steelman, H. R. Gaskins, B. A. C. Harley, Journal of Neuroinflammation 2020, 17, 346.

[15] a)S. C. Woodburn, J. L. Bollinger, E. S. Wohleb, Journal of Neuroinflammation 2021, 18, 258; b)A. Buonfiglioli, D. Hambardzumyan, Acta Neuropathologica Communications 2021, 9, 54.

[16] A. Vidal-Itriago, R. A. W. Radford, J. A. Aramideh, C. Maurel, N. M. Scherer, E. K. Don, A. Lee, R. S. Chung, M. B. Graeber, M. Morsch, Frontiers in Immunology 2022, 13.

[17] L. F. P. Bosch, K. Kierdorf, Frontiers in Cellular Neuroscience 2022, 16.

[18] C.-W. Chang, A. J. Seibel, A. Avendano, M. G. Cortes-Medina, J. W. Song, Advanced Healthcare Materials 2020, 9, 1901399.

[19] S. Quintero-Fabián, R. Arreola, E. Becerril-Villanueva, J. C. Torres-Romero, V. Arana-Argáez, J. Lara-Riegos, M. A. Ramírez-Camacho, M. E. Alvarez-Sánchez, Frontiers in Oncology 2019, 9.

[20] a)C.-W. Chang, H.-C. Shih, M. G. Cortes-Medina, P. E. Beshay, A. Avendano, A. J. Seibel, W.-H. Liao, Y.-C. Tung, J. W. Song, ACS Applied Materials & Interfaces 2023, 15, 15047; b)O.-Y. Hong, H.-Y. Jang, Y.-R. Lee, S. H. Jung, H. J. Youn, J.-S. Kim, Scientific Reports 2022, 12, 12125; c)A. Das, M. Monteiro, A. Barai, S. Kumar, S. Sen, Scientific Reports 2017, 7, 14219; d)S. Zhang, Z. Wan, G. Pavlou, A. X. Zhong, L. Xu, R. D. Kamm, Advanced Functional Materials 2022, 32, 2206767.

[21] a)R. Shi, Y.-Q. Tang, H. Miao, MedComm 2020, 1, 47; b)T. Katopodi, S. Petanidis, D. Anestakis, C. Charalampidis, I. Chatziprodromidou, G. Floros, P. Eskitzis, P. Zarogoulidis, C. Koulouris, C. Sevva, K. Papadopoulos, M. Dagher, V. A. Karakousis, N. Varsamis, V. Theodorou, C. M. Mystakidou, K. Vlassopoulos, S. Kosmidis, N. I. Katsios, K. Farmakis, C. Kosmidis, Frontiers in Immunology 2024, 14.

[22] a)S. H. Baik, S. Kang, W. Lee, H. Choi, S. Chung, J.-I. Kim, I. Mook-Jung, Cell Metabolism 2019, 30, 493; b)J. Cheng, R. Zhang, Z. Xu, Y. Ke, R. Sun, H. Yang, X. Zhang, X. Zhen, L.-T. Zheng, Journal of Neuroinflammation 2021, 18, 129; c)J. P. Bielanin, D. Sun, Translational Stroke Research 2023, 14, 435.

[23] R. Bhargava, Annual Review of Analytical Chemistry 2023, 16, 205.

[24] a)S. J. Blaschke, S. Demir, A. König, J.-A. Abraham, S. U. Vay, M. Rabenstein, D. N. Olschewski, C. Hoffmann, M. Hoffmann, N. Hersch, R. Merkel, B. Hoffmann, M. Schroeter, G. R. Fink, M. A. Rueger, Frontiers in Cellular Neuroscience 2020, 14; b)M. Karlstetter, E. Lippe, Y. Walczak, C. Moehle, A. Aslanidis, M. Mirza, T. Langmann, Journal of Neuroinflammation 2011, 8, 125; c)S. Li, I. Wernersbach, G. S. Harms, M. K. E. Schäfer, Frontiers in Immunology 2022, 13; d)S. F. Spampinato, G. Costantino, S. Merlo, P. L. Canonico, M. A. Sortino, Biomolecules 2022, 12, 1174; e)S. Lively, L. C. Schlichter, Journal of Neuroinflammation 2013, 10, 843.

[25] M. Frühauf, U. Zeitschel, C. Höfling, F. Ullm, F. V. Rabiger, G. Alber, T. Pompe, U. Müller, S. Roßner, European Journal of Neuroscience 2021, 53, 4034.

[26] Y. Liu, J. Zhang, Y. Li, Y. Zhao, S. Kuermanbayi, J. Zhuang, H. Zhang, F. Xu, F. Li, Chemical Science 2024, 15, 171.

[27] A. Procès, Y. A. Alpizar, S. Halliez, B. Brône, F. Saudou, L. Ris, S. Gabriele, Biomaterials 2024, 305, 122426.

[28] a)Y. Yongjun, H. Shuyun, C. Lei, C. Xiangrong, Y. Zhilin, K. Yiquan, J Neuroimmunol 2013, 260, 1; b)D. S. Markovic, K. Vinnakota, N. van Rooijen, J. Kiwit, M. Synowitz, R. Glass, H. Kettenmann, Brain Behav Immun 2011, 25, 624.

[29] M. J. Baker, J. Trevisan, P. Bassan, R. Bhargava, H. J. Butler, K. M. Dorling, P. R. Fielden, S. W. Fogarty, N. J. Fullwood, K. A. Heys, C. Hughes, P. Lasch, P. L. Martin-Hirsch, B. Obinaju, G. D. Sockalingum, J. Sulé-Suso, R. J. Strong, M. J. Walsh, B. R. Wood, P. Gardner, F. L. Martin, Nature Protocols 2014, 9, 1771.

[30] P. R. Griffiths, J. A. d. Haseth, in Fourier Transform Infrared Spectrometry, 2007.

[31] in Fourier Transform Infrared Spectrometry, 2007.

[32] a)N. Shirvaikar, L. A. Marquez-Curtis, A. R. Shaw, A. R. Turner, A. Janowska-Wieczorek, Experimental Hematology 2010, 38, 823; b)H. Raghu, P. K. Sodadasu, R. R. Malla, C. S. Gondi, N. Estes, J. S. Rao, BMC Cancer 2010, 10, 647; c)K. Kajiwara, P.-K. Chen, Y. Abe, S. Okuda, S. Kon, J. Adachi, T. Tomonaga, Y. Fujita, M. Okada, Current Biology 2022, 32, 3460.

[33] a)R. Orihuela, C. A. McPherson, G. J. Harry, British Journal of Pharmacology 2016, 173, 649; b)X. Liu, J. Liu, S. Zhao, H. Zhang, W. Cai, M. Cai, X. Ji, R. K. Leak, Y. Gao, J. Chen, X. Hu, Stroke 2016, 47, 498.

[34] a)M. Naumann, N. Arend, R. R. Guliev, C. Kretzer, I. Rubio, O. Werz, U. Neugebauer, International Journal of Molecular Sciences 2023, 24, 824; b)N. Pavillon, A. J. Hobro, S. Akira, N. I. Smith, Proceedings of the National Academy of Sciences 2018, 115, E2676.

[35] A. J. Boutilier, S. F. Elsawa, International Journal of Molecular Sciences 2021, 22, 6995.

[36] a)J. Renkawitz, A. Kopf, J. Stopp, I. de Vries, M. K. Driscoll, J. Merrin, R. Hauschild, E. S. Welf, G. Danuser, R. Fiolka, M. Sixt, Nature 2019, 568, 546; b)E. D. Tabdanov, N. J. Rodríguez-Merced, A. X. Cartagena-Rivera, V. V. Puram, M. K. Callaway, E. A. Ensminger, E. J. Pomeroy, K. Yamamoto, W. S. Lahr, B. R. Webber, B. S. Moriarity, A. S. Zhovmer, P. P. Provenzano, Nature Communications 2021, 12, 2815.

[37] a)E. F. Willis, S. J. Kim, W. Chen, M. Nyuydzefe, K. P. A. MacDonald, A. Zanin-Zhorov, M. J. Ruitenberg, J. Vukovic, Brain, Behavior, and Immunity 2024, 117, 181; b)A.-E. Roser, L. Tönges, P. Lingor, Frontiers in Aging Neuroscience 2017, 9; c)R. Socodato, A. Rodrigues-Santos, J. Tedim-Moreira, T. O. Almeida, T. Canedo, C. C. Portugal, J. B. Relvas, Cell Death & Disease 2023, 14, 690.

[38] a)G. Cheng, X. Fan, M. Hao, J. Wang, X. Zhou, X. Sun, Molecular Cancer 2016, 15, 30; b)Z. D’Costa, K. Jones, A. Azad, R. van Stiphout, S. Y. Lim, A. L. Gomes, P. Kinchesh, S. C. Smart, W. Gillies McKenna, F. M. Buffa, O. J. Sansom, R. J. Muschel, E. O’Neill, E. Fokas, Cancer Research 2017, 77, 5952; c)O. Prokopchuk, B. Grünwald, U. Nitsche, C. Jäger, O. L. Prokopchuk, E. C. Schubert, H. Friess, M. E. Martignoni, A. Krüger, BMC Cancer 2018, 18, 128.

[39] a)B. Schoeps, J. Frädrich, A. Krüger, Trends in Cell Biology 2023, 33, 413; b)L. Liu, S. Yang, K. Lin, X. Yu, J. Meng, C. Ma, Z. Wu, Y. Hao, N. Chen, Q. Ge, W. Gao, X. Wang, E. W. F. Lam, L. Zhang, F. Li, B. Jin, D. Jin, Scientific Reports 2022, 12, 11181; c)M. Langguth, E. Maranou, S. A. Koskela, O. Elenius, R. E. Kallionpää, E.-M. Birkman, O. I. Pulkkinen, M. Sundvall, M. Salmi, C. R. Figueiredo, Genes & Immunity 2024, 25, 188.

[40] a)P. Ray, A. C. Stacer, J. Fenner, S. P. Cavnar, K. Meguiar, M. Brown, K. E. Luker, G. D. Luker, Oncogene 2015, 34, 2043; b)D. K. Ahirwar, M. W. Nasser, M. M. Ouseph, M. Elbaz, M. C. Cuitiño, R. D. Kladney, S. Varikuti, K. Kaul, A. R. Satoskar, B. Ramaswamy, X. Zhang, M. C. Ostrowski, G. Leone, R. K. Ganju, Oncogene 2018, 37, 4428.

[41] a)N. Goffart, A. Lombard, F. Lallemand, J. Kroonen, J. Nassen, E. Di Valentin, S. Berendsen, M. Dedobbeleer, E. Willems, P. Robe, V. Bours, D. Martin, P. Martinive, P. Maquet, B. Rogister, Neuro-Oncology 2016, 19, 66; b)F. A. Giordano, J. P. Layer, S. Leonardelli, L. L. Friker, R. Turiello, D. Corvino, T. Zeyen, C. Schaub, W. Mueller, E. Sperk, L. C. Schmeel, K. Sahm, S. Kebir, P. Hambsch, T. Pietsch, S. Bisdas, M. Glas, C. Seidel, U. Herrlinger, M. Hölzel, Journal of Clinical Oncology 2023, 41, 2048; c)F. A. Giordano, B. Link, M. Glas, U. Herrlinger, F. Wenz, V. Umansky, J. M. Brown, C. Herskind, Cancers 2019, 11, 272.

[42] Y. Wei, S. Geng, Y. Si, Y. Yang, Q. Chen, S. Huang, X. Chen, W. Xu, Y. Liu, J. Jiang, Advanced Science 2024, 11, 2306715.

[43] M.-C. Liu, H.-C. Shih, J.-G. Wu, T.-W. Weng, C.-Y. Wu, J.-C. Lu, Y.-C. Tung, Lab on a Chip 2013, 13, 1743.

[44] J. Chen, D. J. Laverty, S. Talele, A. Bale, B. L. Carlson, K. A. Porath, K. K. Bakken, D. M. Burgenske, P. A. Decker, R. A. Vaubel, J. E. Eckel-Passow, R. Bhargava, Z. Lou, P. Hamerlik, B. Harley, W. F. Elmquist, Z. D. Nagel, S. K. Gupta, J. N. Sarkaria, Science Translational Medicine 2024, 16, eadj5962.

[45] D. R. Stirling, M. J. Swain-Bowden, A. M. Lucas, A. E. Carpenter, B. A. Cimini, A. Goodman, BMC Bioinformatics 2021, 22, 433.

[46] J. C. Holter, C.-W. Chang, A. Avendano, A. A. Garg, A. K. Verma, M. Charan, D. K. Ahirwar, R. K. Ganju, J. W. Song, Frontiers in Bioengineering and Biotechnology 2022, 10.

[47] a)A. Brand, L. Allen, M. Altman, M. Hlava, J. Scott, Learned Publishing 2015, 28, 151; b)L. Allen, J. Scott, A. Brand, M. Hlava, M. Altman, Nature 2014, 508, 312.

